# The relationship between neural variability and neural oscillations

**DOI:** 10.1101/555649

**Authors:** Edan Daniel, Thomas Meindertsma, Ayelet Arazi, Tobias H. Donner, Ilan Dinstein

**Author notes:** Equal first authorship (Edan Daniel, Thomas Meindertsma).

## Abstract

Neural activity fluctuates over time, creating considerable variability across trials. This trial-by-trial neural variability is dramatically reduced (“quenched”) after the presentation of sensory stimuli. Likewise, the power of neural oscillations, primarily in the alpha-beta band, is also reduced. Despite their similarity, these phenomena have been discussed independently. We hypothesized that the two phenomena are tightly coupled. To test this, we examined magnetoencephalography (MEG) recordings of healthy subjects viewing repeated presentations of a visual stimulus. The timing, amplitude, and spatial topography of variability-quenching and power suppression were remarkably similar. Neural variability quenching was eliminated by excluding the alpha-beta band from the recordings, but not by excluding other frequency-bands. Moreover, individual magnitudes of alpha-beta band power explained 86% of between-subject differences in variability quenching. In contrast, inter-trial-phase-coherence (ITPC) was not correlated with variability quenching. These results reveal that neural variability quenching reflects stimulus-induced changes in the power of alpha-beta band oscillations.

## Introduction

Neural activity is highly variable, such that repeated presentations of an identical stimulus result in variable neural responses across trials (Arieli et al., 1996; Faisal et al., 2008; Shadlen and Newsome, 1998; Tolhurst et al., 1983; Tomko and Crapper, 1974; Werner and Mountcastle, 1963). This trial-by-trial variability is relatively large before stimulus presentation, and strongly reduced (quenched) approximately 200ms after stimulus presentation (Abbott et al., 2011; Arieli et al., 1996; Churchland et al., 2010; He, 2013; Rajan et al., 2010). Neural variability quenching is a robust phenomenon that has been reported in intracellular membrane potential recordings in cats, extracellular recordings of spiking activity in monkeys (Churchland et al., 2010, 2006), and in human electroencephalography (EEG) (Arazi et al., 2017a, 2017b; Schurger et al., 2015), electrocorticography (ECOG) (He and Zempel, 2013), MEG (Schurger et al., 2015), and functional magnetic resonance imaging (fMRI) recordings (Broday-Dvir et al., 2018; He, 2013). Furthermore, the phenomenon was reported during both awake and anaesthetized states, and in several cortical areas (Churchland et al., 2010; He, 2013) using a variety of sensory stimuli (Arazi et al., 2017b; Churchland et al., 2010). Neural variability quenching seems to be a network phenomenon that is apparent across large populations of neighboring neurons regardless of their firing rates or stimulus selectivity (Churchland et al., 2010; Goris et al., 2014).

Another robust phenomenon that is apparent in recordings of electrophysiological mass activity is the reduction of induced oscillatory power approximately 200ms after stimulus presentation. This power suppression predominates in the alpha band (8-13Hz) and is often referred to as event related desynchronization (ERD) (Pfurtscheller and Aranibar, 1977; Pfurtscheller and Lopes da Silva, 1999). It is evident in a spatially selective manner corresponding to the sensory-activated cortical areas (Jensen and Mazaheri, 2010), and coincides with increases in gamma power (>30 Hz) and population spiking activity (Mukamel et al., 2005). It is, therefore, commonly assumed that reductions in alpha power indicate an increase in cortical activity (Neuper et al., 2006).

Quenching of neural variability following stimulus presentation can be driven by two independent mechanisms (Figure 1). First, a stimulus-induced decrease in oscillatory power/amplitude would yield fewer trail-by-trial differences regardless of the precise timing of these oscillations (Figure 1A). Second, a stimulus-evoked increase in phase coherence across trials (i.e., better phase locking across trials) would also yield fewer trial-by-trial differences (Figure 1B). The two mechanisms are not mutually exclusive and may both contribute to the variability quenching phenomenon (Dinstein et al., 2015).

**Figure 1.**
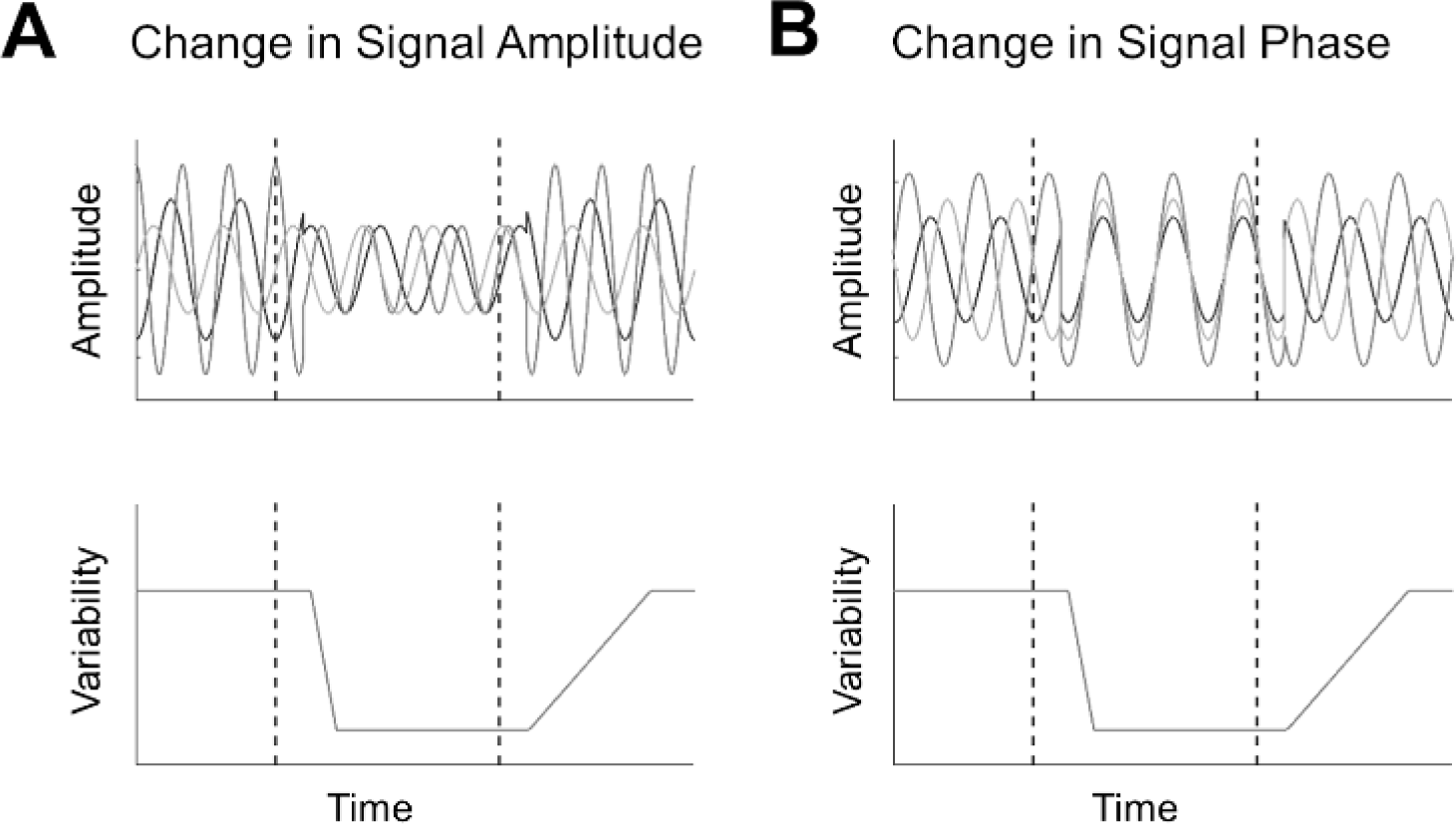
Schematic illustration of two different mechanisms for reducing trial-by-trial variability. Hypothetical oscillatory activity in three independent trials is presented with respect to stimulus presentation (top panels). Dashed lines mark times of stimulus onset and offset, respectively. Reducing the amplitude of oscillations (**A**, top panel) or aligning their phase (**B**, top panel) will create a reduction in trial-by-trial neural variability (bottom panels). Note that the two options are not mutually exclusive.

In the current study we quantified the relationships between spectral power and trial-by-trial neural variability in several ways. First, we extracted specific frequency bands from the MEG data and determined the effect that this had on neural variability magnitudes. Second, we examined whether individual subject differences in spectral power could explain individual differences in neural variability. Finally, we examined whether individual subject differences in inter-trial phase coherence (ITPC) could explain individual differences in neural variability. These analyses were performed using MEG recordings from an experiment with relatively long (750ms), salient, rotating stimuli, because this experimental design was particularly useful for identifying sustained gamma band responses (Meindertsma et al., 2017) that are difficult to identify with other techniques and experimental designs (Whitham et al., 2007; Yuval-Greenberg et al., 2008).

## Materials and Methods

The current study utilized a subset of MEG recordings that were part of a previously published study regarding perceptual decision making (Meindertsma et al., 2017).

### Subjects

23 subjects (13 females; age range, 20-54; mean age, 26.6 years; SD, 7.5 years) were included in the current study. All subjects had normal or corrected-to-normal vision and no known history of neurological disorders. The experiment was conducted in accordance with the Declaration of Helsinki and approved by the local ethics committee of the Hamburg Medical Association. Each subject gave written informed consent.

### Experimental design

Subjects passively viewed a repeating visual stimulus while MEG data was recorded (Figure 2A). The stimulus consisted of a large, full-field grid of white crosses (17° X 17°) that rotated in the clock-wise or counter-clockwise direction (speed: 160°/s). This moving stimulus surrounded a full contrast Gabor patch (diameter, 2°; two cycles), located in the lower right or left visual field quadrant (counterbalanced between subjects) and modulated at a temporal frequency of 10 Hz. Subjects fixated on a fixation mark (red outline, white inside, 0.8° width and length) in the middle of the screen. Stimuli were presented using the Presentation Software (NeuroBehavioral Systems Inc.). Stimuli were back-projected on a transparent screen using a Sanyo PLC-XP51 projector with a resolution of 1024X768 pixels at 60 Hz. Subjects were seated 58 cm from the screen in a whole-head magnetoencephalography (MEG) scanner setup in a dimly lit room. Each trial started with the presentation of the fixation mark (750-1250ms), followed by presentation of the full stimulus (750ms), fixation mark (750ms), and an inter-trial-interval of 750ms containing a blank screen. This experiment was used as a localizer for quantifying sensory responses to the rotating mask and Gabor stimuli in a previous study (Meindertsma et al., 2017). This previous study examined perceptual decision making during motion induced blindness (Bonneh et al., 2001), a phenomenon where the moving stimulus (mask) induces the illusory disappearances of small but salient static stimuli (i.e., the Gabor). The flicker of the Gabor stimulus in the localizer, however, was specifically implemented to prevent the occurrence of motion induced blindness (Meindertsma et al., 2017).

**Figure 2.**
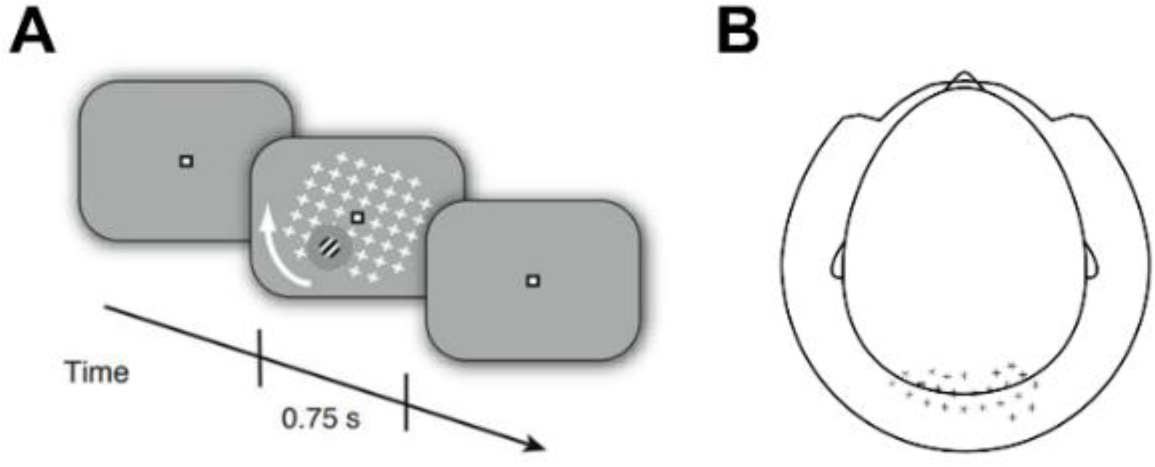
Experimental design. **A,** Illustration of the stimulus presented to the subjects. A rotating mask of white crosses was presented along with a peripheral Gabor patch located in either the left of right bottom quadrants of the visual field. Each trial began with a fixation mark (750-1250ms), followed by the stimulus (750ms), another fixation mark period (750ms), and finally an inter-trial interval with a blank screen (750ms). **B,** Scalp map indicating the location of chosen sensors that were used in subsequent analyses.

### Data acquisition

MEG data were acquired using a 275-channel MEG system (VSM/CTF Systems) with a sample rate of 1200 Hz, while subjects were in a seated position. The location of each subjects’ head was measured throughout the experiment using three fiducial markers placed on both ears and the nasal bridge to control for excessive movement. Furthermore, electrooculography and electrocardiography were recorded to aid in post-hoc artifact rejection.

### Preprocessing

MEG data were analyzed in MATLAB (MathWorks Inc., USA) using the Fieldtrip toolbox (Oostenveld et al., 2011), EEG toolbox (Delorme and Makeig, 2004a), and custom-written software. Each trial was defined as an epoch that started 800ms before stimulus onset and lasted until 1000ms after stimulus offset (i.e., −800 to 1750ms with respect to stimulus onset). We detected artifacts related to environmental noise, eye and muscle activity, and squid jumps using standard automated artifact rejection methods included in the Fieldtrip toolbox. Trials containing artifacts were excluded and remaining data was down sampled to 500 Hz. The final analysis was conducted, on average, with 156 trials (SD = 54.6) per subject. We focused our analysis on the cortical regions processing the physical stimulus (i.e., visual cortex). We, therefore, selected 25 occipital sensors that exhibited the strongest stimulus-induced response, as defined and previously reported by Meindertsma et al. (2017) (Figure 2B). One sensor was missing in many subjects, and therefore removed from the data of all subjects, resulting in 24 sensors of interest. We also present topographical displays of our findings, which demonstrate the spatial selectivity of the results.

### Spectral analyses

Spectral decomposition of MEG recordings was performed using a sliding Hamming-window Fourier transform (step size: 40ms, window length: 500ms), as implemented in EEGLAB (Delorme and Makeig, 2004b), and performed separately for each trial, sensor, and subject. Power was calculated for each time-frequency segment by computing the absolute values of the Fourier coefficients. The resulting time–frequency power estimates (i.e., spectrograms), aligned to stimulus onsets, were averaged across the 24 sensors described above and then across trials to obtain a spectrogram for each subject. We then isolated power changes in specific frequency bands, which included the delta (1-4 Hz), alpha-beta (5-25 Hz), and gamma (60-120 Hz) bands. Topographic plots of power were obtained by isolating a time-window of interest and averaging the power across this window, per sensor, across subjects.

Relative stimulus-induced change in power was normalized into units of percent signal change. We calculated the power in the pre-stimulus (*Power*_*pre*_) period (−250ms to stimulus onset) and post-stimulus (*Power*_*post*_) period (200ms to 700ms after stimulus onset) and computed percent signal change as follows:

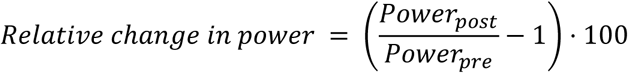

Finally, we also estimated inter-trial phase coherence (ITPC) across trials, for each frequency and sensor (Delorme and Makeig, 2004a). This measure reflects the degree to which the phase of each frequency is aligned across trials. ITPC values were then normalized to percentage change units with respect to the pre-stimulus baseline as described above for the power calculations.

### Neural Variability Analyses

Trial-by-trial variability was computed across trials for each time point in every sensor. Absolute trial-by-trial variability in the pre-stimulus (*Var*_*pre*_) and post-stimulus (*Var*_*post*_) periods were computed by averaging across the relevant time-points (−250ms to stimulus onset, and 200ms to 700ms after stimulus onset, respectively). Relative change in trial-by-trial variability (i.e., neural variability quenching) was then estimated by dividing the variability in the post-stimulus period by the pre-stimulus period and adjusting to percentage change units, as follows:

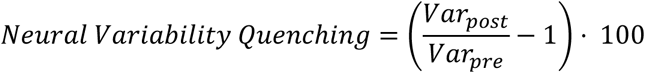

To isolate the contribution of each frequency band to the magnitude of variability quenching, we used Hamming windowed finite impulse response filters to isolate or exclude data in specific frequency bands. We then compared the magnitude of variability quenching before and after applying each filter to the data. This included band-pass filters to isolate the data in delta (1-4Hz), alpha-beta (4-25Hz), or gamma (60-120Hz) bands as well as band-stop filters that excluded the data of each of these frequency band.

### Head motion

To control for excessive motion, we computed the three-dimensional position of the head in every time-point. We then quantified head-motion by computing the mean absolute difference in position from each time-point to the next. Estimated magnitudes of head-motion were then correlated with individual measures of neural variability quenching to determine potential relationships or lack there-of.

### Statistical tests

To identify statistically significant changes in oscillatory power or trial-by-trial variability, while correcting for multiple comparisons, we used two-tailed cluster-based permutation tests (Efron and Tibshirani, 1994). This involved identifying time-points with an un-corrected p-value smaller than 0.05 when applying a paired sample t-test. Consecutive time-points that exceeded the threshold formed candidate clusters and the sum of each cluster’s t-values was computed. We then used a Monte-Carlo permutation with 1000 iterations to define a probability distribution of t-value sums from clusters in randomly drawn sets of time-points (Efron and Tibshirani, 1994). The corrected p-value was defined by the relative percentile of the actual cluster statistic relative to this null distribution of random cluster statistics.

We computed Pearson’s correlation coefficients to assess potential relationships between neural variability quenching and oscillatory power or ITPC, across subjects. The same analysis was performed in the control analysis with estimates of head motion. We also used a partial-correlation analysis to estimate the relative contribution of oscillatory power in each frequency band to the magnitude of neural variability. This eliminated inter-dependencies across frequency bands, thereby isolating the contribution of each frequency band from that of the others.

## Results

Subjects exhibited strong stimulus-induced responses (Figure 3), with a characteristic and well-known time-frequency and spatial signature (Donner and Siegel, 2011). An initial broadband power increase in all frequencies was followed by different frequency-band specific dynamics. Power in the delta (1-4Hz) band increased dramatically with stimulus presentation, peaking at ~+200ms, and remained significantly larger than the pre-stimulus period throughout stimulus presentation. Power in the alpha-beta frequency band (5-25Hz) increased transiently and then decreased to negative values ~200ms after stimulus presentation (Figure3B). Power in the gamma (60-120Hz) band increased in a sustained manner after stimulus presentation and returned to pre-stimulus levels ~250ms after stimulus offset.

**Figure 3.**
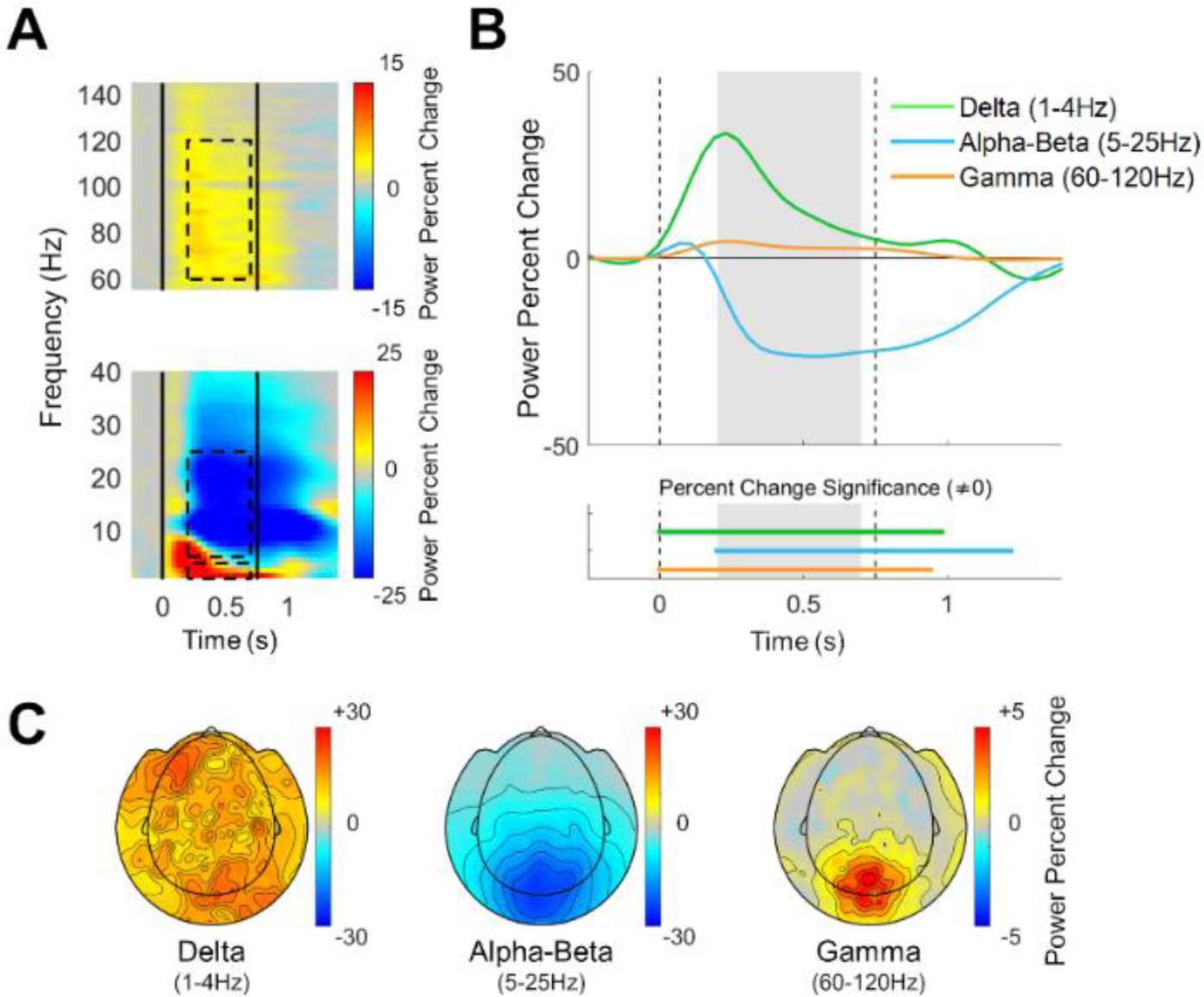
Stimulus-induced responses in the different frequency bands. **A,** Spectrogram demonstrating the relative change in power with respect to the pre-stimulus period (−250ms to stimulus onset). Black vertical lines: stimulus onset and offset. Dashed rectangles: selected frequency bands and time window. **B,** Temporal changes in the power of each frequency band, averaged across the selected sensors, all trials, and subjects. Dashed vertical lines: stimulus onset and offset. Gray filling: window over which neural variability quenching was computed in subsequent analyses. Horizontal lines on bottom indicate time segments where the change in power of each band was significantly different from zero (p < 0.05, two-tailed permutation test, cluster corrected). **C,** Topographic maps of mean power change 200-700ms after stimulus presentation, relative to the pre-stimulus period in units of percent signal change (averaged across trials and subjects).

Note the similar temporal dynamics of power across the alpha-beta frequency range (Figure 3A), which justify its selection as a single band, as also used in other recent studies (Michalareas et al., 2016). Power changes in the three selected frequency bands, 200-700ms after stimulus presentation, exhibited different spatial characteristics (Figure 3C). Power reduction in the alpha-beta band and power enhancement in the gamma band were specific to sensors located over occipital and parietal cortices, while changes in the delta band were not.

### Neural variability quenching

Subjects also exhibited robust reductions in trial-by-trial neural variability 200-700ms after stimulus presentation in comparison to the pre-stimulus period (Figure 4A). To determine the relationship between variability quenching and the activity of specific frequency bands, we re-computed neural variability after isolating each frequency band using band-pass filters (Figure 4B). This revealed that variability quenching was remarkably strong 200-700ms after stimulus presentation, in isolated alpha-beta band activity, where variability quenching reached a mean value of −53.3% (blue line, Figure 4B). In contrast, neural variability quenching was absent in isolated delta band activity (−0.3%, green line, Figure 4B) and neural variability was enhanced, rather than quenched in isolated gamma band activity (+7%, orange line, Figure 4B). Note that variability quenching in the original, un-filtered data reached a mean value of −20.3% (yellow line, Figure 4B).

**Figure 4.**
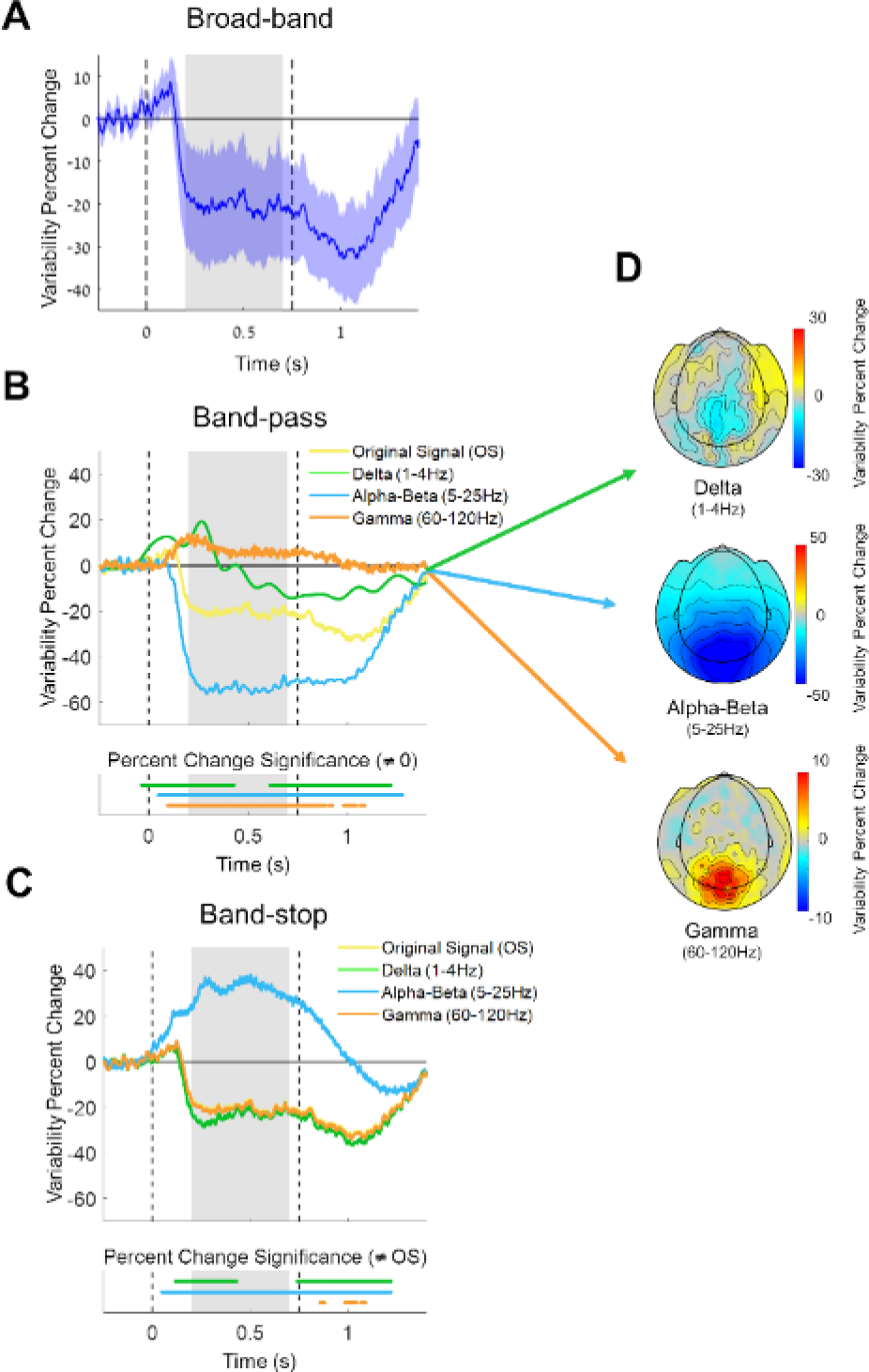
Trial-by-trial neural variability changes following stimulus presentation. **A,** Neural variability over time in units of percent change relative to pre-stimulus period (averaged across selected sensors and all subjects). Blue shaded area: confidence interval across subjects. Dashed vertical lines: stimulus onset and offset. Gray background: window with sustained neural variability quenching. **B**, Neural variability quenching within each isolated frequency band (delta, alpha-beta, or gamma) when using band-pass filters. Time segments where variability was significantly different from zero are marked in the lower panel (p < 0.05, two-tailed permutation test, cluster corrected). **C,** Neural variability quenching after eliminating each frequency band from the data using band-stop filters. Time segments where variability was significantly different from that in the original signal are marked in the lower panel (p < 0.05, two-tailed permutation test, cluster corrected). **D**, Topographic maps of neural variability changes 200-700ms after stimulus presentation, relative to the pre-stimulus period, after isolating each of the frequency bands.

In a complementary analysis we used band-stop filters to eliminate specific frequency bands in the MEG data (see Materials and Methods) and then re-computed trial-to-trial variability (Figure 4C). Eliminating the alpha-beta frequency band dramatically altered variability quenching from a mean value of −20.3% in the original signal (yellow line, Figure 4C) to variability enhancement with a mean value of +32.4% (blue line, Figure 4C). In contrast, eliminating the Delta or Gamma bands had minor effects on variability quenching, which had values of −23.6% and −20.8% (green and orange lines, Figure 4C) respectively. Taken together, these results demonstrate that variability quenching is mostly driven by neural activity changes in the alpha-beta band. Note that this analysis estimated the mean response across subjects and disregarded individual differences.

Differences in the spatial topography of neural variability were also apparent when isolating each of the frequency bands (Figure 4D). Neural variability changes were diffused and patchy in isolated delta band activity. In contrast, neural variability changes in a spatially selective manner (i.e., in occipital and parietal sensors) in isolated alpha-beta band activity and gamma band activity. The band-specific spatial changes in neural variability (Figure 4D) were very similar to the spatial changes in power (Figure 3C). This was apparent in a moderate spatial correlation in the delta (r(268) = 0.5, p=0.03) band and very strong correlations in the alpha-beta (r(268) = 0.95, p<0.001) and gamma (r(268) = 0.95, p<0.001) bands.

### Individual differences

In line with previous studies (Arazi et al., 2017b), individual subjects exhibited distinct magnitudes of neural variability quenching. These individual differences were strongly correlated with the magnitudes of stimulus-induced power changes (Figure 5) in the delta (r(23) = 0.62, p=0.002) and alpha-beta (r(23) = 0.93, p<0.001) frequency bands, but not in the gamma band (r(23) = −0.17, p=0.44). These results demonstrate that individual differences in variability quenching were best explained by individual differences in alpha-beta band power reductions, which single-handedly explained the vast majority of differences across subjects (r squared = 0.86). To examine the combined predictive value of power changes in all three frequency bands on variability quenching magnitudes, we also performed a multiple regression analysis. The regression model included three predictors containing the individual subjects’ power changes in each frequency band. This regression model yielded an adjusted r squared value of 0.88, suggesting that adding the delta and gamma power changes did very little to improve the ability to predict individual variability quenching magnitudes.

**Figure 5.**
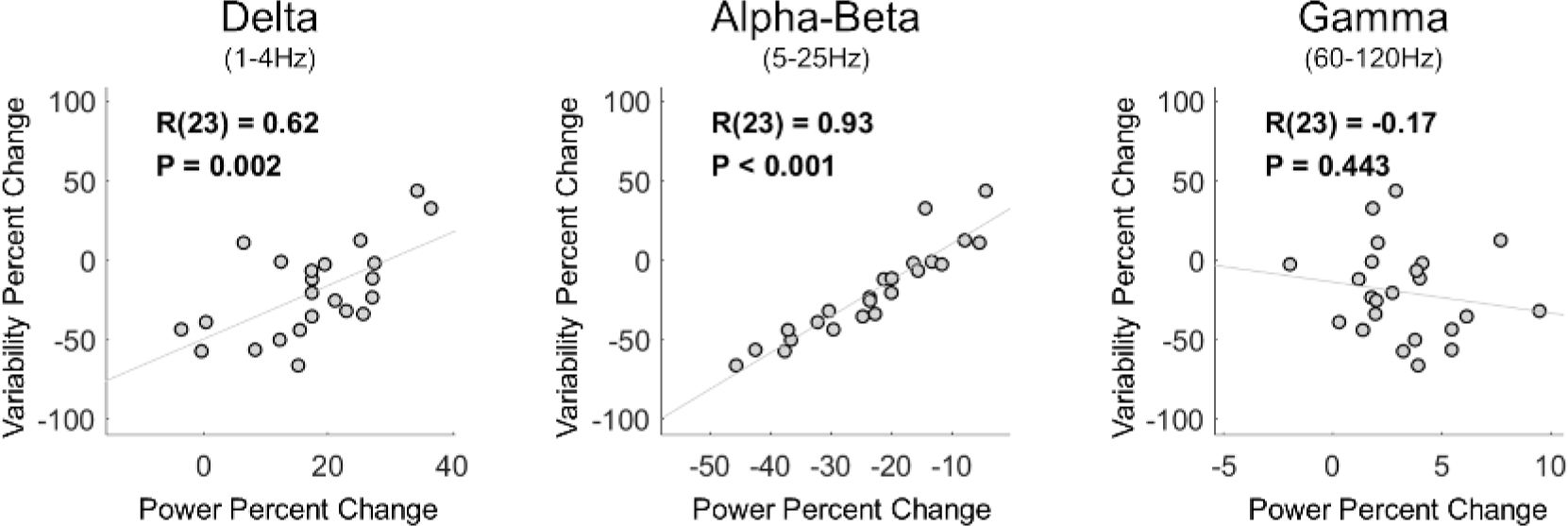
Individual magnitudes of variability-quenching were strongly correlated with relative changes in MEG power of specific frequency bands. Scatter plots demonstrating the correlation between individual magnitudes of variability quenching and changes in delta, alpha-beta, or gamma band power. Linear fit lines, Pearson's correlation coefficients (R) and significance level (P) are presented in each panel.

### Variability quenching is not associated with the timing of neural responses

The results presented thus far establish that stimulus-induced modulations in alpha-beta band oscillatory power contribute to neural variability quenching. Trial-to-trial variability, however, is governed not only by the amplitude of neural oscillations (Figure 1A), but also by their phase (i.e., timing) relative to stimulus presentation (Figure 1B). We, therefore, tested whether variability quenching was also associated with an increase in inter trial phase coherence (ITPC). ITPC increased transiently after stimulus onset and offset with a time course and spectral profile (Figure 6A) that resembled broadband changes in power (Figure 3A). This ITPC time course was distinct from that of variability quenching, which was sustained throughout the stimulus presentation (Figure 4). Furthermore, individual magnitudes of variability quenching were not significantly correlated with ITPC changes in any of the examined frequency bands: delta (r(23) = −0.05, p=0.83), alpha-beta (r(23) = −0.30, p=0.17), or gamma (r(23) = −0.03, p=0.91) (Figure 6B). Taken together, these results demonstrate that variability quenching is not associated with stimulus-evoked changes in the phase of neural oscillations.

**Figure 6.**
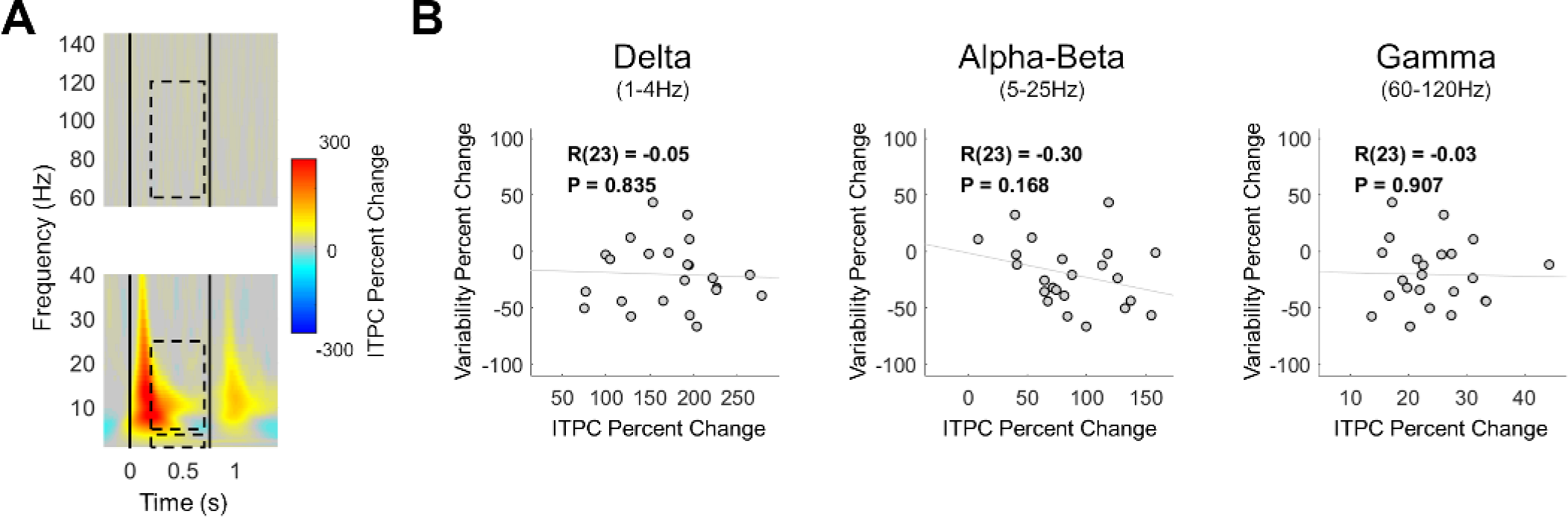
Inter-trial phase coherence (ITPC) was not associated with neural variability quenching. **A,** Time/frequency representation of stimulus evoked changes in ITPC, relative to the pre-stimulus baseline, revealed transient rather than sustained changes. Black vertical lines: times of stimulus onset and offset. Dashed rectangles: selected frequency bands and time window (for comparison with the previous analyses). **B,** Scatter plots demonstrating the lack of correlations between changes in band specific ITPC and changes in neural variability, for the delta, alpha-beta, or gamma frequency bands. Linear fit lines, Pearson's correlation coefficients (R) and significance level (P) are presented in each panel.

### Control analyses

To exclude alternative explanations of the data we examined whether head motion affected our measures of neural variability. We computed a mean measure of head motion magnitude for each subject (see Methods) and found that there was no significant correlation with neural variability quenching (r(23) = 0.12, p=0.54). Between-subject differences in variability quenching were, therefore, unrelated to individual differences in head movements.

## Discussion

Our results suggest that neural variability quenching after stimulus presentation reflects stimulus-induced changes in the amplitude of alpha-beta band oscillations. The timing, amplitude, and spatial topography of neural variability quenching and decreases in oscillatory power in the alpha-beta band were remarkably similar (Figure 3&4). Indeed, removing the alpha-beta band from the data eliminated the neural variability quenching phenomenon (Figure 4C). The strong relationship between changes in oscillatory power in this frequency band and neural variability were also apparent when examining individual subject differences (Figure 5). Specifically, individual magnitudes of alpha-beta band power changes explained 86% of between-subject variability in neural variability quenching. In contrast, power changes in the delta or gamma bands had little effect on the variability quenching phenomenon (Figure 4). While individual differences in delta band power were correlated with individual magnitudes of variability quenching (Figure 5), including the delta band differences in a multiple regression model together with the alpha-beta band differences had a negligible effect on explanatory power (adjusted r squared = 0.88). Gamma band differences were not correlated with variability quenching magnitudes. Taken together, these findings suggest that neural variability quenching is a product of specific changes in alpha-beta band oscillatory power that are induced by the presentation of a stimulus.

While neural variability quenching was strongly related to the amplitude of induced oscillatory power, it was not related to the phase-locking of neural oscillations. The time course of ITPC changes did not correspond to the time course of neural variability quenching, and individual subject ITPC magnitudes were not correlated with neural variability quenching magnitudes (Figure 6). Taken together, these results suggest that cortical responses to sensory stimuli are characterized by relatively high reproducibility (i.e., low trial-by-trial variability) that is driven by stimulus-induced decreases in alpha-beta band oscillations rather than stimulus-evoked phase resetting.

### Event related desynchronization and synchronization

Stimulus and/or task induced decreases in oscillatory power, primarily in the alpha-beta band, are commonly referred to as ERD (Pfurtscheller and Aranibar, 1977; Pfurtscheller and Lopes da Silva, 1999), which is thought to coincide with increased synchronization in gamma band oscillations and increased multi-unit activity (Mukamel et al., 2005). These concomitant changes in cortical oscillatory power are thought to represent the transition of cortical state from an idle “resting-state” to an active state of anticipation, sensory processing, and/or task initiation. It is believed that such oscillatory changes are essential for synchronizing the activity of task-related cortical neural ensembles (Engel et al., 2001; Siegel et al., 2012). Previous studies have not examined how these changes in oscillatory power relate to measures of trial-by-trial neural variability.

We selected the current experiment, utilizing MEG recordings and an experimental design with relatively long (750ms), salient, rotating stimuli, because this experimental setup is particularly useful for identifying sustained gamma band responses (Meindertsma et al., 2017) that are difficult to identify with other techniques and experimental designs (Whitham et al., 2007; Yuval-Greenberg et al., 2008). Indeed, our results revealed clear sustained gamma band responses with focal topography in occipital and parietal sensors (Figure 3) and differences in the amplitude of gamma power across subjects (Figure 5).

We initially hypothesized that we would find a positive correlation between the magnitude of ERD and neural variability quenching as well as a negative correlation between the magnitude of gamma band power and neural variability quenching. This was expected given the inverse relationship between stimulus-induced ERD and gamma synchronization. Despite the strong relationship between neural variability quenching and ERD, we did not find a significant relationship between gamma synchronization and neural variability quenching (Figure 5). This suggests that individual differences in trial-by-trial neural variability are mostly governed by differences in alpha-beta band oscillations that are far larger in amplitude than gamma band oscillations.

### Ongoing neural activity and stimulus evoked/induced responses

Ongoing neural activity continuously changes and fluctuates in the absence of stimuli or tasks thereby creating considerable moment-by-moment neural variability (Arieli et al., 1996; Biswal et al., 1995; Fox and Raichle, 2007). Some studies have suggested that these ongoing fluctuations persist during the processing of stimuli (Arieli et al., 1996) and execution of tasks (Becker et al., 2011; Fox et al., 2007) such that stimulus evoked responses are linearly superimposed on ongoing fluctuations. In such a case one would expect similar trial-by-trial neural variability to exist before and after stimulus presentation. Many recent studies, however, have shown that trial-by-trial neural variability is dramatically reduced following stimulus presentation (Arazi et al., 2017a, 2017b; Broday-Dvir et al., 2018; Churchland et al., 2010; Goris et al., 2014; He, 2013; Schurger et al., 2015). This suggests that ongoing neural fluctuations do not persist, but are instead altered by the presentation of a stimulus such that trial-by-trial variability is reduced. This alteration could be in the form of a decrease in induced oscillatory power and/or an increase in phase coherence (Figure 1) (Dinstein et al., 2015).

Our results suggest that the presentation of a visual stimulus quenches trial-by-trial neural variability by reducing induced oscillatory power in the alpha-beta band, rather than evoking a reproducible phase locked response (Figures 3-6). The strong relationship between the ERD and the variability quenching phenomena suggests that ongoing neural activity fluctuations are actively suppressed after the presentation of a sensory stimulus, perhaps to achieve a more reproducible and stable cortical state during sensory processing (Schurger et al., 2015).

### Behavioral significance

There are several similarities in the behavioral significance that has been assigned to the ERD and neural variability phenomena. For example, some have reported that allocating attention reduces neural variability across trials (Broday-Dvir et al., 2018; Cohen and Maunsell, 2009; Mitchell et al., 2009, 2007) while others have reported that allocating attention creates ERD (Ikkai et al., 2016; Siegel et al., 2008; Thut, 2006). Similarly, some have reported that threshold-level stimuli are accurately perceived on trials with reduced neural variability (Schurger et al., 2015, 2010) while others have reported the same on trials with larger ERD (van Dijk et al., 2008). Finally, individuals with lower contrast discrimination thresholds exhibited larger magnitudes of neural variability quenching, which coincided with larger ERD magnitudes (Arazi et al., 2017a). All studies, except the last one, have reported one measure or the other and none have directly compared ERD and neural variability measures. We speculate that these independent studies may be reporting strongly correlated measures that seem to describe a common underlying neural mechanism with specific behavioral effects.

### Conclusions

Our results suggest that stimulus-induced reductions in alpha-beta band power cause the observed reduction in trial-by-trial variability of broadband signals. The suppression of these oscillations, and subsequent reduction in trial-by-trial variability, may enable sensory cortices to generate more stable and reproducible neural representations of a stimulus across trials, which is likely to be beneficial for accurate perception. This appealing conceptual framework, whereby cortical responses involve an active reduction of neural “noise” (i.e., neural activity that is not related to the stimulus), is in line with signal detection theory principles (Green and Swets, 1966), and fits well with the existing literature regarding neural variability quenching and ERD. Quantifying oscillatory power, inter-trial phase coherence, and trial-by-trial variability in the same experiments will enable future studies to assess the robustness and validity of this conceptual framework across different stimuli, tasks, and recording techniques, and determine its behavioral significance.

## Funding Information

This work was supported by the Israeli Science Foundation (Grant 961/14 to I.D.), the Netherlands Organization for Scientific Research (NWO, 406-14-016 to T.H.D. and T.M.) and the German Research Foundation (DFG, Heisenberg Professorship DO 1240/3-1 and DO 1240/4-1 to T.H.D.).

## References

Abbott LF, Rajan K, Sompolinsky H. 2011. Interactions between Intrinsic and Stimulus-Evoked Activity in Recurrent Neural Networks, The Dynamic Brain: An Exploration of Neuronal Variability and Its Functional Significance. doi:10.1093/acprof:oso/9780195393798.003.0004

Arazi A, Censor N, Dinstein I. 2017a. Neural Variability Quenching Predicts Individual Perceptual Abilities. J Neurosci 37:97–109. doi:10.1523/JNEUROSCI.1671-16.2016

Arazi A, Gonen-Yaacovi G, Dinstein I. 2017b. The Magnitude of Trial-By-Trial Neural Variability Is Reproducible over Time and across Tasks in Humans. eNeuro 4:ENEURO.0292--17.2017. doi:10.1523/ENEURO.0292-17.2017

Arieli A, Sterkin A, Grinvald A, Aertsen A. 1996. Dynamics of Ongoing Activity : Explanation of the Large Variability in Evoked Cortical Responses 273:1868–1871.

Becker R, Reinacher M, Freyer F, Villringer A, Ritter P. 2011. How ongoing neuronal oscillations account for evoked fMRI variability. J Neurosci 31:11016–27. doi:10.1523/JNEUROSCI.0210-11.2011

Biswal B, Yetkin FZ, Haughton VM, Hyde JS. 1995. Functional Connectivity in the Motor Cortex of Resting Human Brain Using Echo-Planar Mri. Magn Reson Med. doi:DOI 10.1002/mrm.1910340409

Bonneh YS, Cooperman A, Sagi D. 2001. Motion-induced blindness in normal observers. Nature 411:798–801. doi:10.1038/35081073

Broday-Dvir R, Grossman S, Furman-Haran E, Malach R. 2018. Quenching of spontaneous fluctuations by attention in human visual cortex. Neuroimage 171:84–98. doi:10.1016/j.neuroimage.2017.12.089

Churchland MM, Yu BM, Cunningham JP, Sugrue LP, Cohen MR, Corrado GS, Newsome WT, Clark AM, Hosseini P, Scott BB, Bradley DC, Smith MA, Kohn A, Movshon JA, Armstrong KM, Moore T, Chang SW, Snyder LH, Lisberger SG, Priebe NJ, Finn IM, Ferster D, Ryu SI, Santhanam G, Sahani M, Shenoy K V. 2010. Stimulus onset quenches neural variability: A widespread cortical phenomenon. Nat Neurosci 13:369–378. doi:10.1038/nn.2501

Churchland MM, Yu BM, Ryu SI, Santhanam G, Shenoy K V. 2006. Neural Variability in Premotor Cortex Provides a Signature of Motor Preparation. J Neurosci 26:3697–3712. doi:10.1523/JNEUROSCI.3762-05.2006

Cohen MR, Maunsell JHR. 2009. Attention improves performance primarily by reducing interneuronal correlations. Nat Neurosci. doi:10.1038/nn.2439

Delorme A, Makeig S. 2004a. EEGLAB: An open source toolbox for analysis of single-trial EEG dynamics including independent component analysis. J Neurosci Methods 134:9–21. doi:10.1016/j.jneumeth.2003.10.009

Delorme A, Makeig S. 2004b. EEGLAB: an open source toolbox for analysis of single-trial EEG dynamics including independent component analysis. J Neurosci Methods 134:9–21. doi:10.1016/j.jneumeth.2003.10.009

Dinstein I, Heeger DJ, Behrmann M. 2015. Neural variability: friend or foe? Trends Cogn Sci 19:322–8. doi:10.1016/j.tics.2015.04.005

Donner TH, Siegel M. 2011. A framework for local cortical oscillation patterns. Trends Cogn Sci. doi:10.1016/j.tics.2011.03.007

Efron B, Tibshirani R. 1994. An introduction to the bootstrap. Chapman and Hall/CRC.

Engel AK, Fries P, Singer W. 2001. Dynamic predictions: Oscillations and synchrony in top–down processing. Nat Rev Neurosci. doi:10.1038/35094565

Faisal A, Selen L, Wolpert D. 2008. Noise in the nervous system. Nat Rev Neurosci 9:292–303.

Fox MD, Raichle ME. 2007. Spontaneous fluctuations in brain activity observed with functional magnetic resonance imaging. Nat Rev Neurosci 8:700–11. doi:10.1038/nrn2201

Fox MD, Snyder AZ, Vincent JL, Raichle ME. 2007. Intrinsic Fluctuations within Cortical Systems Account for Intertrial Variability in Human Behavior. Neuron. doi:10.1016/j.neuron.2007.08.023

Goris RLT, Movshon JA, Simoncelli EP. 2014. Partitioning neuronal variability. Nat Neurosci 17:858–865. doi:10.1038/nn.3711

Green DM, Swets JA. 1966. Signal detection theory and psychophysics. Society. doi:10.1901/jeab.1969.12-475

He BJ. 2013. Spontaneous and Task-Evoked Brain Activity Negatively Interact. J Neurosci 33:4672–4682. doi:10.1523/JNEUROSCI.2922-12.2013

He BJ, Zempel JM. 2013. Average Is Optimal: An Inverted-U Relationship between Trial-to-Trial Brain Activity and Behavioral Performance. PLoS Comput Biol 9. doi:10.1371/journal.pcbi.1003348

Ikkai A, Dandekar S, Curtis CE. 2016. Lateralization in Alpha-Band Oscillations Predicts the Locus and Spatial Distribution of Attention. PLoS One 11:e0154796. doi:10.1371/journal.pone.0154796

Jensen O, Mazaheri A. 2010. Shaping Functional Architecture by Oscillatory Alpha Activity: Gating by Inhibition. Front Hum Neurosci 4:186. doi:10.3389/fnhum.2010.00186

Meindertsma T, Kloosterman NA, Nolte G, Engel AK, Donner TH. 2017. Multiple Transient Signals in Human Visual Cortex Associated with an Elementary Decision. J Neurosci 37:5744–5757. doi:10.1523/JNEUROSCI.3835-16.2017

Michalareas G, Vezoli J, van Pelt S, Schoffelen J-M, Kennedy H, Fries P. 2016. Alpha-Beta and Gamma Rhythms Subserve Feedback and Feedforward Influences among Human Visual Cortical Areas. Neuron 89:384–397. doi:10.1016/j.neuron.2015.12.018

Mitchell JF, Sundberg KA, Reynolds JH. 2009. Spatial Attention Decorrelates Intrinsic Activity Fluctuations in Macaque Area V4. Neuron. doi:10.1016/j.neuron.2009.09.013

Mitchell JF, Sundberg KA, Reynolds JH. 2007. Differential Attention-Dependent Response Modulation across Cell Classes in Macaque Visual Area V4. Neuron. doi:10.1016/j.neuron.2007.06.018

Mukamel R, Gelbard H, Arieli A, Hasson U, Fried I, Malach R. 2005. Coupling between neuronal firing, field potentials, and fMRI in human auditory cortex. Science (80-) 309:951–954. doi:10.1126/science.1110913

Neuper C, Wörtz M, Pfurtscheller G. 2006. ERD/ERS patterns reflecting sensorimotor activation and deactivation. Prog Brain Res. doi:10.1016/S0079-6123(06)59014-4

Oostenveld R, Fries P, Maris E, Schoffelen JM. 2011. FieldTrip: Open source software for advanced analysis of MEG, EEG, and invasive electrophysiological data. Comput Intell Neurosci 1:1–9. doi:10.1155/2011/156869

Pfurtscheller G, Aranibar A. 1977. Event-related cortical desynchronization detected by power measurements of scalp EEG. Electroencephalogr Clin Neurophysiol 42:817–826. doi:10.1016/0013-4694(77)90235-8

Pfurtscheller G, Lopes da Silva FH. 1999. Event-related EEG/MEG synchronization and desynchronization: basic principles. Clin Neurophysiol 110:1842–1857. doi:10.1016/S1388-2457(99)00141-8

Rajan K, Abbott LF, Sompolinsky H. 2010. Stimulus-dependent suppression of chaos in recurrent neural networks. Phys Rev E-Stat Nonlinear, Soft Matter Phys 82:1–5. doi:10.1103/PhysRevE.82.011903

Schurger A, Pereira F, Treisman A, Cohen JD. 2010. Reproducibility distinguishes conscious from nonconscious neural representations. Science 327:97–99. doi:10.1126/science.1180029

Schurger A, Sarigiannidis I, Naccache L, Sitt JD, Dehaene S. 2015. Cortical activity is more stable when sensory stimuli are consciously perceived. Proc Natl Acad Sci 112:E2083––E2092. doi:10.1073/pnas.1418730112

Shadlen MN, Newsome WT. 1998. The variable discharge of cortical neurons: implications for connectivity, computation, and information coding. J Neurosci 18:3870–3896. doi:0270–6474/98/183870-27$05.00/0

Siegel M, Donner TH, Engel AK. 2012. Spectral fingerprints of large-scale neuronal interactions. Nat Rev Neurosci 13:121–34. doi:10.1038/nrn3137

Siegel M, Donner TH, Oostenveld R, Fries P, Engel AK. 2008. Neuronal Synchronization along the Dorsal Visual Pathway Reflects the Focus of Spatial Attention. Neuron 60:709–719. doi:10.1016/j.neuron.2008.09.010

Thut G. 2006. -Band Electroencephalographic Activity over Occipital Cortex Indexes Visuospatial Attention Bias and Predicts Visual Target Detection. J Neurosci 26:9494–9502. doi:10.1523/JNEUROSCI.0875-06.2006

Tolhurst DJ, Movshon JA, Dean AF. 1983. The statistical reliability of single neurons in cat and monkey visual cortex. Vision Res 23:775–785. doi:10.1016/0042-6989(83)90200-6

Tomko GJ, Crapper DR. 1974. Neuronal variability: non-stationary responses to identical visual stimuli. Brain Res 79:405–418. doi:10.1016/0006-8993(74)90438-7

van Dijk H, Schoffelen J-M, Oostenveld R, Jensen O. 2008. Prestimulus Oscillatory Activity in the Alpha Band Predicts Visual Discrimination Ability. J Neurosci. doi:10.1523/JNEUROSCI.1853-07.2008

Werner G, Mountcastle VB. 1963. the Variability of Central Neural Activity in a Sensory System, and Its Implications for the Central Reflection of Sensory Events. J Neurophysiol 26:958–977. doi:10.1152/jn.1963.26.6.958

Whitham EM, Pope KJ, Fitzgibbon SP, Lewis T, Clark CR, Loveless S, Broberg M, Wallace A, DeLosAngeles D, Lillie P, Hardy A, Fronsko R, Pulbrook A, Willoughby JO. 2007. Scalp electrical recording during paralysis: Quantitative evidence that EEG frequencies above 20 Hz are contaminated by EMG. Clin Neurophysiol 118:1877–1888. doi:10.1016/j.clinph.2007.04.027

Yuval-Greenberg S, Tomer O, Keren AS, Nelken I, Deouell LY. 2008. Transient Induced Gamma-Band Response in EEG as a Manifestation of Miniature Saccades. Neuron 58:429–441. doi:10.1016/j.neuron.2008.03.027

